# Mesoporous silica particles functionalized with newly extracted fish oil (*Omeg@Silica*) inhibit lung cancer cell growth

**DOI:** 10.1101/2021.04.20.440579

**Authors:** Caterina Di Sano, Claudia D’Anna, Antonino Scurria, Claudia Lino, Mario Pagliaro, Rosaria Ciriminna, Elisabetta Pace

**Affiliations:** Istituto per la Ricerca e l’Innovazione Biomedica, CNR, Via U. La Malfa 153, 90146 Palermo, Italy; Istituto per lo Studio dei Materiali Nanostrutturati, CNR, Via U. La Malfa 153, 90146 Palermo, Italy

**Keywords:** omega-3, lung cancer, cancer cells, cancer growth, fish oil, PUFA

## Abstract

*Omeg@Silica* microparticles consisting of whole fish oil rich in omega-3 lipids, vitamin D_3_ and zeaxanthin extracted with biobased limonene from anchovy fillet leftovers (*AnchoisOil*) encapsulated within mesoporous silica particles are highly effective in modulating oxidative stress, mitochondrial damage or in promoting antitumor effects in lung cancer cells. A panel of three different human non-small cell lung cancer (NSCLC) cell lines (A549, Colo 699 and SKMES) was used. Cancer cells were treated with *AnchoisOil* dispersed in ethanol (10 and 15 μg/ml) or encapsulated in silica, and cell cycle, reactive oxigen species (ROS) and mitochondrial stress (MitoSOX) assessed by flow cytometry. The effects on long-term proliferation (clonogenic assay) were also evaluated. The sub-micron *Omeg@Silica* microparticles were more effective than fish oil alone in increasing ROS and mitocondrial damage, in altering cell cycle as well as in reducing colony formation ability in the tested lung cancer cell lines. These results suggest that *Omeg@Silica* mesoporous silica functionalized with whole fish oil has antitumor effects in NSCLC cell lines and support its investigation in lung cancer therapy.

## 1. Introduction

Lung cancer is the leading form of cancer worldwide in terms of incidence and death rate,^1^ with non-small cell lung cancer (NSCLC) accounting for 85% of lung cancer cell types. Despite advances in detection and improvements to standard of care, NSCLC is often diagnosed at an advanced stage and bears poor prognosis. There is a widespread, urgent and global need to implement therapies to improve prognosis of NSCLC.

Regular intake of omega-3 (*n*-3) marine lipids rich in long-chain polyunsaturated fatty acids (PUFAs) such as docosahexaenoic acid (DHA, C22:6*n*-3) and eicosapentaenoic acid (EPA, C20:5*n*-3) is recommended by most world’s national and international health authorities for the prevention of many chronic diseases, including cancer.^2^ Following associative epidemiological evidence (changing with time) originating from Dyerberg and Bang discovery of very low incidence of ischemic heart-disease and complete absence of diabetes mellitus in the 1960s Greenland’s Inuit population,^3^ detailed physiological, pharmacological and pathophysiological evidence has given clear insights into the benefit from balancing the *n*-6 with *n*-3 lipid molecular mediators in the arachidonic acid cascade.^4^

A plethora of studies have demonstrated that ω-3 PUFAs exert therapeutic role against certain types of cancer,^5^ including lung cancer.^6^ These essential lipids exert inhibitory effects on lung cancer growth by reducing tumor cell proliferation or by increasing cell apoptosis.^7^ In 2014, Newman and co-workers discovered that in lung carcinoma cell lines, fish oil derived EPA reduces non‐small cell lung cancer cellular proliferation through prostaglandin E2 (PGE_2_) formation by cyclooxygenase-2 (COX-2) enzymes and downregulation of phosphoinositide 3-kinase (PI3K) pathways in NSCLC cells.^8^ Subsequent investigation on the inhibitory role of DHA on NSCLC *in vitro* and on fat-1 transgenic mice identified in resolvin D1, an eicosanoid metabolite of DHA, the molecule responsible to significantly inhibit lung cancer cell growth and invasion by increasing expression of miR-138-5p microRNA precursor,^9^ with concomitant results suggesting that miR-138-5p is indeed a tumor suppressor in NSCLC cells.^10^ Preclinical studies have shown evidence also that omega-3 PUFA metabolites modulate pivotal pathways underlying complications secondary to cancer, up-regulating anti-inflammatory lipid mediators such as protectins, maresins, and resolvins.^11^

Unfortunately, a nearly global dietary unbalance between *n*-3 and *n*-6 essential lipids emerged during the 10^th^ and 20^th^ centuries which has led to the present very low to low range of blood EPA + DHA for most of the world’s population (6.5 out of 7 billion people in 2015) increasing global risk for chronic disease.^12^

The omega-3 lipid deficit in the diet of most industrialized countries has led to the rapid development of omega-3 dietary supplement industry based on highly refined fish oil, with a significant impact on increasing overfishing across the oceans and in the Mediterranean sea.^13^ These trends originate two urgent needs: to develop sustainable sources of marine lipids,^14^ and to create formulations that may potentiate and expand the physiological activity of formulations based on marine oils, for example avoiding refinement which eliminates from fish oil the lipophilic natural phlorotannins which crucially protect fish oil omega-3 from oxidation and autoxidation.^15^

The present study reports the *in vivo* anticancer effects of a new formulation (*Omeg@Silica*) comprised of an integral fish oil (*AnchoisOil*) rich in omega-3 lipids, vitamin D3 and zeaxanthin extracted with biobased limonene from anchovy fillet leftovers^16^ encapsulated within mesoporous silica particles (see Figure S1).^17^ Experiments were carried out testing both the newly sourced *AnchoisOil* whole fish oil and *Omeg@Silica* microparticles suspended in aqueous ethanol (FOS) in limiting lung cancer cell growth using a panel of lung cancer cell lines (A549, Colo699 and SK-MES-1).

## 2. Materials and Methods

### Extraction of AnchoisOil

Fish oil was easily obtained from anchovy fillet leftovers kindly donated by an anchovy fillet company based in Sicily (Agostino Recca Conserve Alimentari Srl, Sciacca, Italy) according to a published procedure introduced in 2019.^16^ In detail, 204 g of frozen anchovy waste in the blender jar of an electric blender was added with a first aliquot of 106 g of *d*-limonene (Acros Organics, 96%) refrigerated at 4 °C. After grinding twice for 15 s each time to mix and homogenize the anchovy leftovers, a semi-solid grey mixture was obtained which was extracted with limonene, an high boling citrus-derived natural solvent with many health-beneficial properties.^18^ An aliquot (50.7 g) of this mixture was transferred in a glass beaker and added with 51.4 g of limonene.

The solid-liquid extraction proceeded by magnetically stirring at 700 rpm the mixture kept at room temperature in the beaker closed with an aluminum foil and further sealed with parafilm. After 21 h, the yellow supernatant obtained was transferred to the evaporating balloon of a rotary evaporator (Büchi Rotovapor R-200 equipped with a V-700 vacuum pump and V-850 vacuum controller) to remove the solvent under reduced pressure (40 mbar) at 90 °C. An oily extract weighing 3.0 g deeply colored in orange (*AnchoisOil*) remained in the evaporating balloon after limonene evaporation. The oil contains a good amount (81.5 μg/kg) of vitamin D in the form of bioactive isomer vitamin D_3_ (cholecalciferol), in good agreement with the typical amounts of vitamin D_3_ found in fish oils.^19^ The limonene solvent almost entirely recovered thanks to the recirculation chiller supporting the rotary evaporator with sufficient cooling to condense the vaporized solvent can be used in a subsequent extraction run.

### Preparation of Omeg@Silica

The *Omeg@Silica* particles used throughout this study are MCM-41 silica particles loaded with 50 wt% sustainably sourced anchovy fish oil whose highly reproducible synthesis was lately described.^17^ In detail, the synthesis of the MCM-41 material was carried out according to a published template-assisted published sol-gel process.^20^ After dissolving 1 g of hexadecyltrimethylammonium bromide (CTAB ≥99% pure, Aldrich) and 280 mg of sodium hydroxide (Analyticals, Carlo Erba) in 480 mL of deionized water, an aliquot (5.4 mL) of tetraethylorthosilicate (TEOS ≥99% pure, Aldrich) was added dropwise to the solution. The resulting mixture was kept under continuous mechanical stirring (400 rpm) at 80 °C for 2 h. The solid precipitate was recovered by filtration, washed with abundant deionized water and methanol (99.8% pure, Aldrich) and dried at 60°C for 48 h. The residual surfactant entrapped in the silicate was removed via calcination in air at 550°C in a furnace for 6 h.

The resulting MCM-41 mesoporous silica kept in a glass flask under mild mechanical agitation material was added dropwise with the *AnchoisOil*. Addition of a first 60 μL aliquot of fish oil to FMCM-41 (100 mg), was followed by subsequent additions of 20 μL aliquots. After 8 min, addition of oil was complete and a material with a 50% w/w fish oil load was left under agitation for 24 h. Figure S1 displays the resulting *Omeg@Silica* next to the free AnchoisOil.

The material has a low (0.3) polydispersity index and a large negative (−37.6 mV) value of the zeta potential, both important characteristics to develop stable drug delivery applications,^21^ particularly in aqueous phase as the negatively charged particles repel each other preventing aggregation with the dispersion remaining electrostatically stable. The Z-average size (measure of the average size of the particle size distribution resulting by dynamic light scattering) indicates a moderate increase of the average particle size from 217 to 269 nm going from empty mesoporous silica to silica filled with 50 wt% fish oil.

### Preparation of FOS

The aqueous formulations of different concentrations used throughout this study were obtained by proper dilution of mother suspensions of *Omeg@Silica*, *AnchoisOil* and MCM-41 in 10 v/v% ethanol/ PBS Dulbecco’s Phosphate Buffered Saline (Euroclone, Pero, Italy). The *Omeg@Silica* mother suspension was prepared adding 10.1 mg of material to 10 mL of 10 v/v% ethanol/ PBS solution prepared by adding 1 mL of ethanol to 9 mL of PBS. The *AnchoisOil* and the MCM-41 mother suspensions had a concentration of the single components equivalent to that of *Omeg@Silica* mother suspension. In particular, 5 mg of *AnchoisOil* were suspended in 10 mL of 10 v/v% ethanol/ PBS and 5.1 mg of MCM-41 were suspended in 10 mL of 10 v/v% ethanol/ PBS.

### Cell cultures

In this study were used three different human non-small cell lung cancer (NSCLC) cell lines, A549, Colo 699 and SK-MES-1. Eagle’s minimum essential medium, supported with 10% heat‐deactivated (56°C, 30□min) fetal bovine serum (FBS), 1% nonessential amino acids, 2□mM L‐glutamine and 0.5% gentamicin were used for SK-MES-1. RPMI‐1640 medium supplemented with heat‐deactivated (56°C, 30□min) 10% FBS, streptomycin and penicillin, 1% nonessential amino acids and 2□mM L‐glutamine (all from Euroclone) was used for culturing A549 and Colo 699.The cells were maintained in an incubator at 37□°C with a humidified atmosphere with 5% CO_2_and were maintained as adherent monolayers. The cells were grown grown in polystyrene flasks 25 cm^2^ (BD Falcon, Franklin Lakes, New Jersey) to 90% confluence and passaged by trypsin/EDTA.

### Treatment of the cells

Cells were seeded on six-well plates and were cultured to confluence; then the serum was reduced from 10% to 1% in the medium and the cells were treated with two different concentrations (5, 10 and 15 μg/ml) of AnchoisOildispersed in ethanol, silica sub-micron particles and AnchoisOil encapsulated in silica. At the end of stimulation, cells were collected for further evaluations. At least four replicates were performed for each experiment.

### Cell viability/metabolic assay

To assess the right concentration of the stimuli to add to the culture, we used theCellTiter 96^®^ AQueous One Solution Cell Proliferation Assay(PROMEGA, Madison WI USA), a colorimetric method for determining the number of viable cells in proliferation, cytotoxicity or chemosensitivity assays. Cells were plated in 96-well plates and were treated for 24 h in triplicate with of AnchoisOildispersed in ethanol (fish oil) (5, 10 and 15 μg/ml), silica sub-micron particles (5, 10 and 15 μg/ml) and AnchoisOil encapsulated in silica (FOS) (5, 10 and 15 μg/ml). Then 20mL of One Solution reagent contains MTS [3-(4,5-dimethylthiazol-2-yl)-5-(3-carboxymethoxyphenyl)-2-(4-sulfopheyl)2H-tetrazolium]was added to each well, and incubated for 20 min at 37°C, 5% CO2. The absorbance was read at 490 nm on the Microplate reader WallacVictor2 1420 Multilabel Counter (Perkin Elmer). The results were calculated as the percentage of absorbance with respect to that of the control (untreated cells=NT).

### Clonogenic assay

Clonogenic assay was performed as previously described (Pace E et al). In six-well plates, a lower layer was prepared using complete medium in 0.5% agarose. The plates were stored at 4°C for 24 hours. The cell lines were stimulated in 1% FBS medium with the different treatments, as previous described, for 24 hours; then they were harvested and seeded (5 × 10^4^) on the upper layer with 0.3% agarose prepared with the same medium as the lower layer, and finally incubated for 21 days at 37 °C in an atmosphere containing 0.5% CO_2_. At the end of incubation, colonies were counted under an inverted phase-contrast microscope (Leitz, Wetzlar, Germany). The experiment was conducted in triplicate. Colonies were defined as cell aggregates with at least 40 cells. Results are expressed as percentage of non-treated control (NT). All experiments were performed in triplicate.

### Cell Cycle

The cell lines were stimulated in 1% FBS medium with the different treatments, as previous described, for 24 hours. After harvesting, the cells were washed twice with ice-cold PBS and resuspended at 1 × 10^6^ cells/ml in hypotonic fluorochrome solution (0.1% sodium citrate, 0.03% Nonidet P-40 and 50 μg/ml propidium iodide) for 30 min at room temperature in the dark. Then, the cells were acquired and analyzed by flow cytometry. On the basis of DNA content apoptotic cells as M1 (sub-G1 phase), G0/G1 cells as M2, S cells as M3 and G2/M cells as M4 were identified. Data were expressed as percentage of cells.

### Evaluation of mitochondrial stress

The mitochondrial stress was evaluated by the MitoSOX Kit (Molecular Probes Waltham, MA, USA). The cell lines were stimulated in 1% FBS medium with the different treatments, as previous described, for 3 hours. After stimulation cells were collected and, after the addition of the MitoSOX reagent at the concentration 3 μM, they were incubated for 15 min at 37 °C. At the end of the incubation the cells were washed twice in PBS 1% FBS, followed by flow cytometric analysis. Data were expressed as percentage of cells.

### Evaluation of intracellular reactive oxygen species (ROS)

Intracellular reactive oxygen species (ROS) were measured by the conversion of the non-fluorescent dichlorofluorescein diacetate (DCFH-DA; Sigma) in a highly fluorescent compound, DCF, by monitoring the cellular esterase activity in the presence of peroxides, as previously described (Bruno et al. 2011). The cell lines were stimulated in 1% FBS medium with the different treatments, as previous described, for 3 hours. After stimulation cells, ROS generation was assessed by the uptake of 1 μM DCFH-DA, incubation for 10 min at room temperature in the dark. At the end of the incubation, the cells were washed twice in PBS and acquired on a FACSCalibur™ flow cytometer, supported by CellQuest acquisition and data analysis software (Becton Dickinson, Mountain View, CA, USA). Data were expressed as percentage of cells.

### Statistics

Data were expressed as mean ± SD and analysed by Paired t-test. A p-value of less than 0.05 was considered as statistically significant.

## 3. Results

### 3.1 Effects of fish oil, silica and FOS on cell viability /metabolic activity in NSCLC cell lines

Initially, the effects of fish oil (5, 10 and 15 μg/ml), silica (5, 10 and 15 μg/ml) and fish oil in silica (FOS) (5, 10 and 15 μg/ml) on cell viability/metabolic activity after 24 hours were assessed in NSCLC cell lines (A549, Colo699 and SKMES). In A549, silica at 5, 10 and 15 μg/ml and FOS at 10 and 15 μg/ml were able to reduce cell viability /metabolic activity when compared to control and FOS at 10 μg/ml FOS exerted a significant stronger activity on cell viability/metabolic activity than fish oil (Figure 1 A). In Colo699, only FOS at 10 μg/ml was able to reduce cell viability /metabolic activity when compared to control (Figure 1 B). In SKMES, neither the silica nor oil nor FOS at all the tested concentrations significantly modified cell viability /metabolic activity (Figure 1 C).

**Figure 1.**
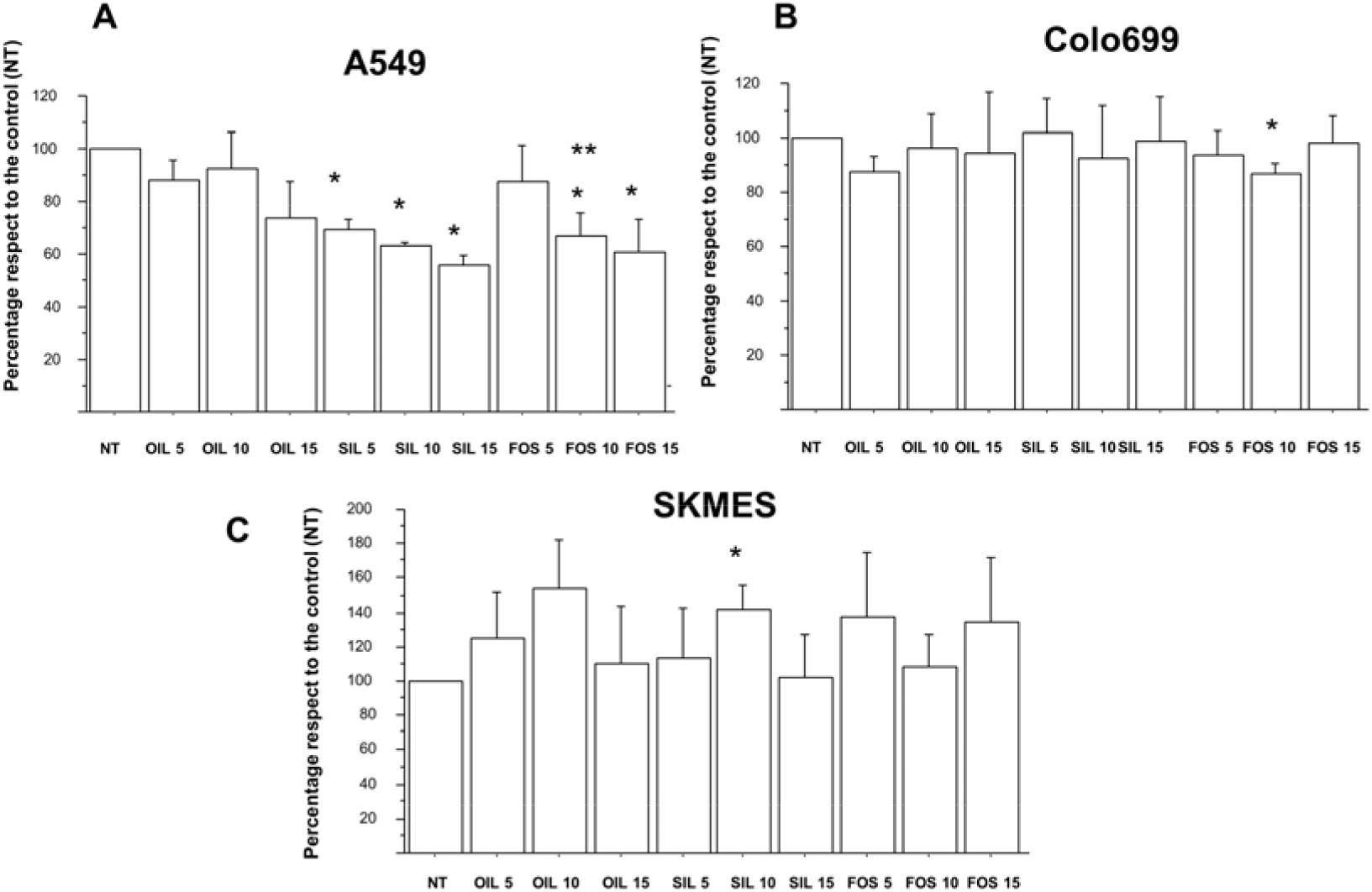
Effect of fish oil, silica and FOS on cell viability /metabolic activity in NSCLC cell lines. NSCLC cell lines, A549 (**A**), Colo699 (**B**), and SKMES (**C**) were cultured for 24 hours with fish oil (5, 10 and 15 μg/ml), silica (5, 10 and 15 μg/ml) and fish oil in silica (FOS) (5, 10 and 15 μg/ml) and cell viability/metabolic activity was assessed by MTS. Data are expressed as % of NT and represent mean ± S.D. (n=3). * p<0.05

### 3.2 Effects of fish oil, silica and FOS on cell cycle events in NSCLC cell lines

The effects of fish oil alone (10 and 15 μg/ml), silica (10 and 15 μg/ml) and FOS (10 and 15 μg/ml) in cell cycle of the NSCLC cell lines (A549, Colo699 and SKMES) were tested. In A549, fish oil alone at both the tested concentrations was able to significantly reduce M3 (fish oil at 10 and at 15 μg/ml *vs.* NT: p= 0.0129 and p= p=0.02) without any effects on M1, M2 or M4. Silica at 10 μg/ml but not at 15 μg/ml was able to increase M1 (silica at 10 μg/ml *vs.* NT: p=0.014) and Silica at 15 μg/ml was able to significantly reduce M3 (silica 15 μg/ml *vs.* NT: p= 0.006).

FOS at 10 μg/ml was able to significantly increase M1 (FOS at 10 μg/ml *vs.* NT: p=0.019) without any effects on M2, M3 or M4. This effect was significantly higher than the effects mediated by fish oil alone at the same concentration (FOS at 10 μg/ml *vs.* fish oil at 10 μg/ml p=0.014) (Figures 2A-2B). In Colo699, fish oil alone at both the tested concentrations was able to significantly reduce M3 (*a*: fish oil at 10 and 15 μg/ml *vs.* NT: p=0.0012 and p=0.005) without any effects on M1 and M2. Fish oil at 10 μg/ml significantly reduced M4 (fish oil at 10 μg/ml vs NT: p=0.007.

**Figure 2.**
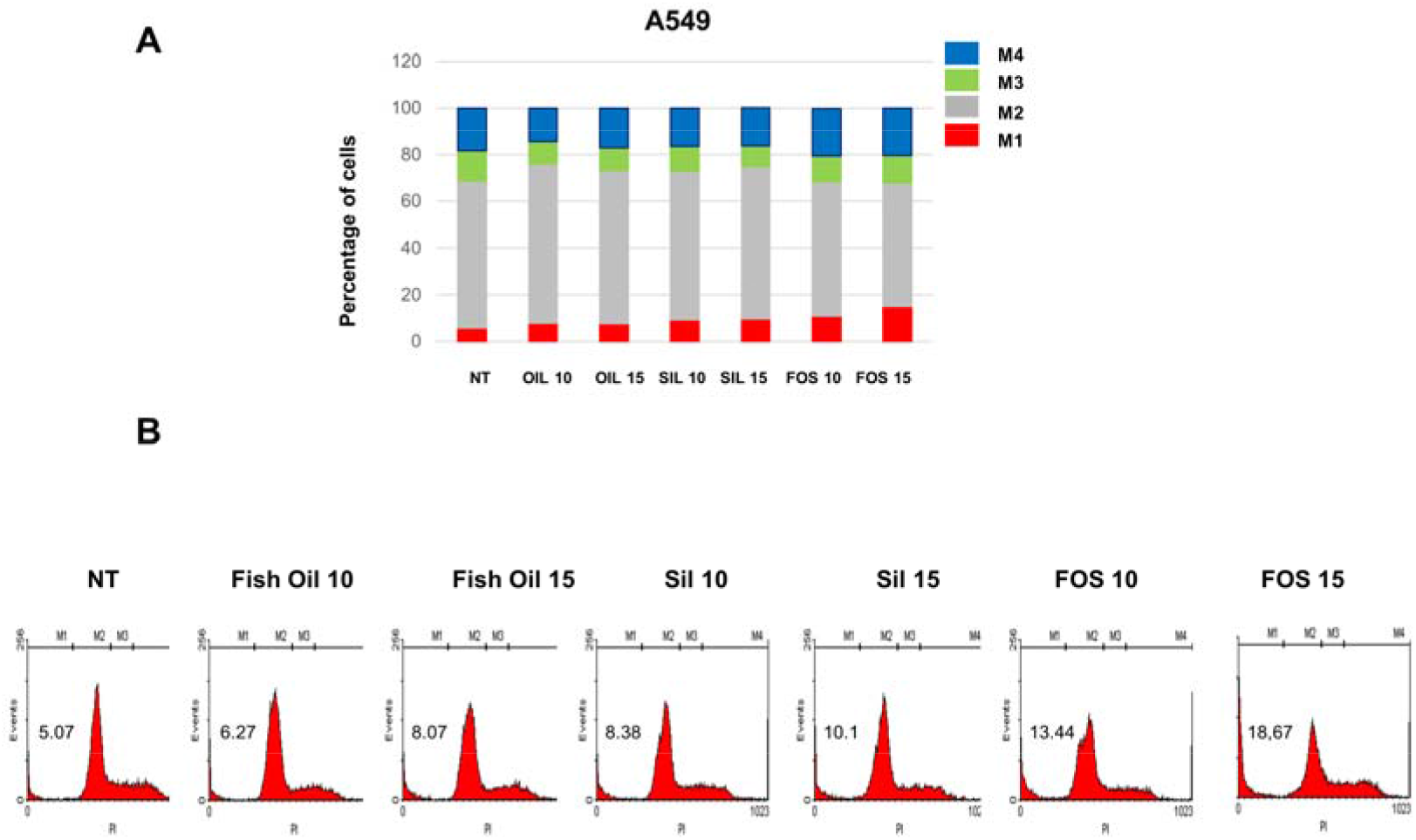
Effect of fish oil, silica and FOS on cell cycle events in A549 cells. A549 were cultured for 24 hours with fish oil (10 and 15 μg/ml), silica (10 and 15 μg/ml) and fish oil in silica (FOS) (10 and 15 μg/ml) and cell cycle was assessed by flow cytometry. **A.** Data are expressed as percentage of cells (mean±SD). * p<0.05. **B.** Representative histograms were shown.

Periodic mesoporous silica at 15 μg/ml significantly reduced M4 (*b*: silica at 15 μg/ml *vs.* NT: p<0.003) without any effects on M1, M2 and M3.

FOS at 10 μg/ml was able to significantly increase M1 (FOS at 10 μg/ml vs NT: p=0.008) and this effect was significantly higher than the effects mediated by fish oil alone at the same concentration (*c*: FOS at 10 μg/ml vs fish oil at 10 μg/ml p=0.037). FOS at 10 μg/ml significantly reduced M4 (FOS at 10 μg/ml vs NT: p=0.025) without any effects on M2 and M3 (Figures 3A-3B).

**Figure 3.**
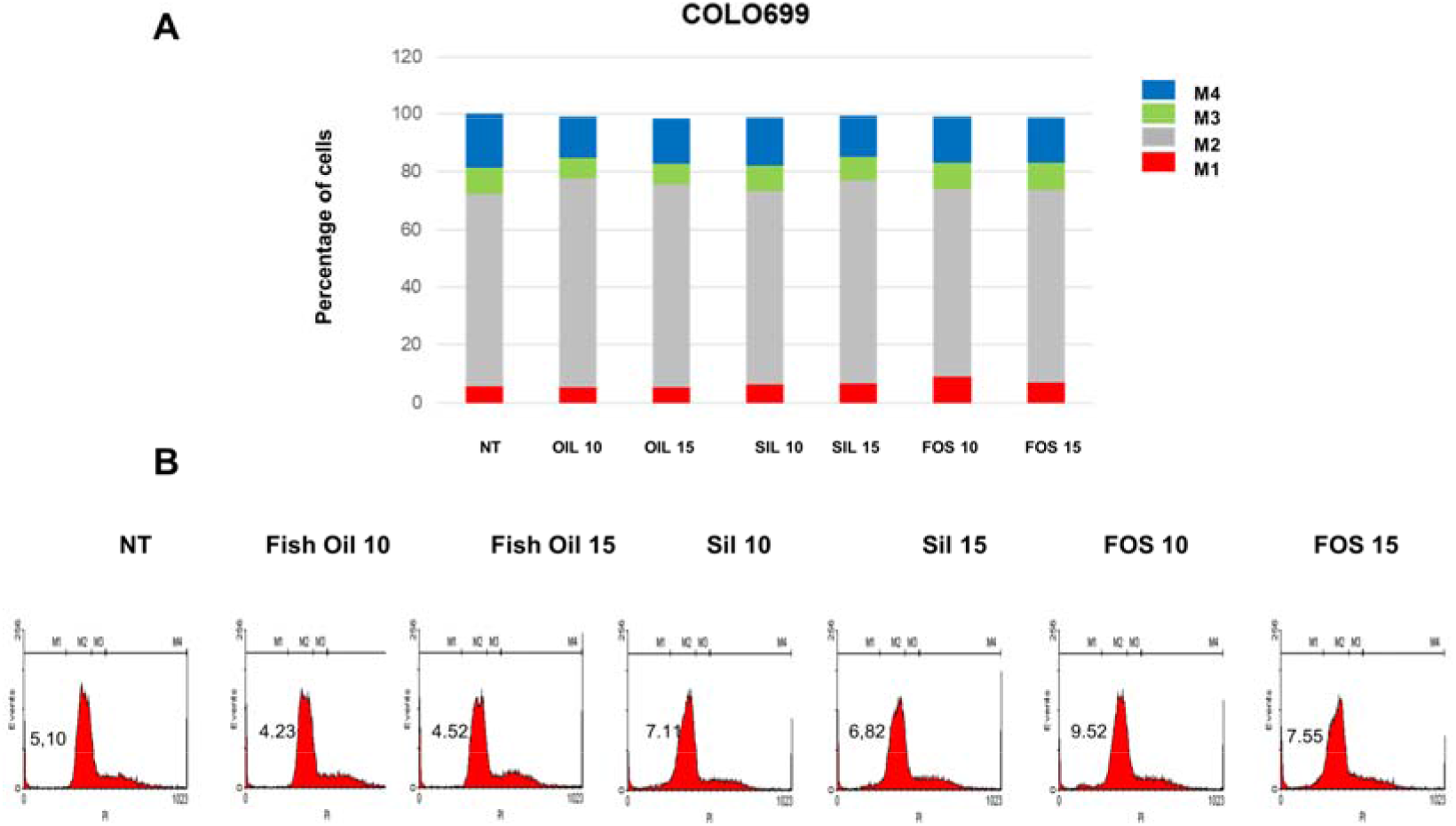
Effect of fish oil, silica and FOS on cell cycle events in Colo699. Colo699 were cultured for 24 hours with fish oil (10 and 15 μg/ml), silica (10 and 15 μg/ml) and fish oil in silica (FOS) (10 and 15 μg/ml) and cell cycle was assessed by flow cytometry. **A.** Data expressed as percentage of cells (mean±SD). * p<0.05. **B.** Representative histograms are shown.

In SKMES: fish oil alone at both the tested concentrations was able to significantly reduce M3 (*a*: fish oil at 10 and at 15 μg/ml *vs.* NT: p= 0.005 and p=0.001) and M4 (fish oil at 10 and at 15 μg/ml vs NT: p= 0.02 and p=0.003) while only at 10 μg/ml was able to increase M2 (fish oil at 10 μg/ml *vs.* NT: p=0.03); silica at 15 μg/ml significantly increased M2 and reduced M3 (*b*: silica at 15 μg/ml *vs.* NT: p=0.002); FOS at 10 μg/ml significantly increased M2 (FOS at 10 μg/ml *vs.* NT: p=0.01) and reduced M3 (FOS at 10 μg/ml *vs.* NT: p=0.0009) while FOS at both the tested concentrations significantly reduced M4 (*c*: FOS at 10 and at 15 μg/ml *vs.* NT: p= 0.03 and p=0.03) (Figures 4A-4B).

**Figure 4.**
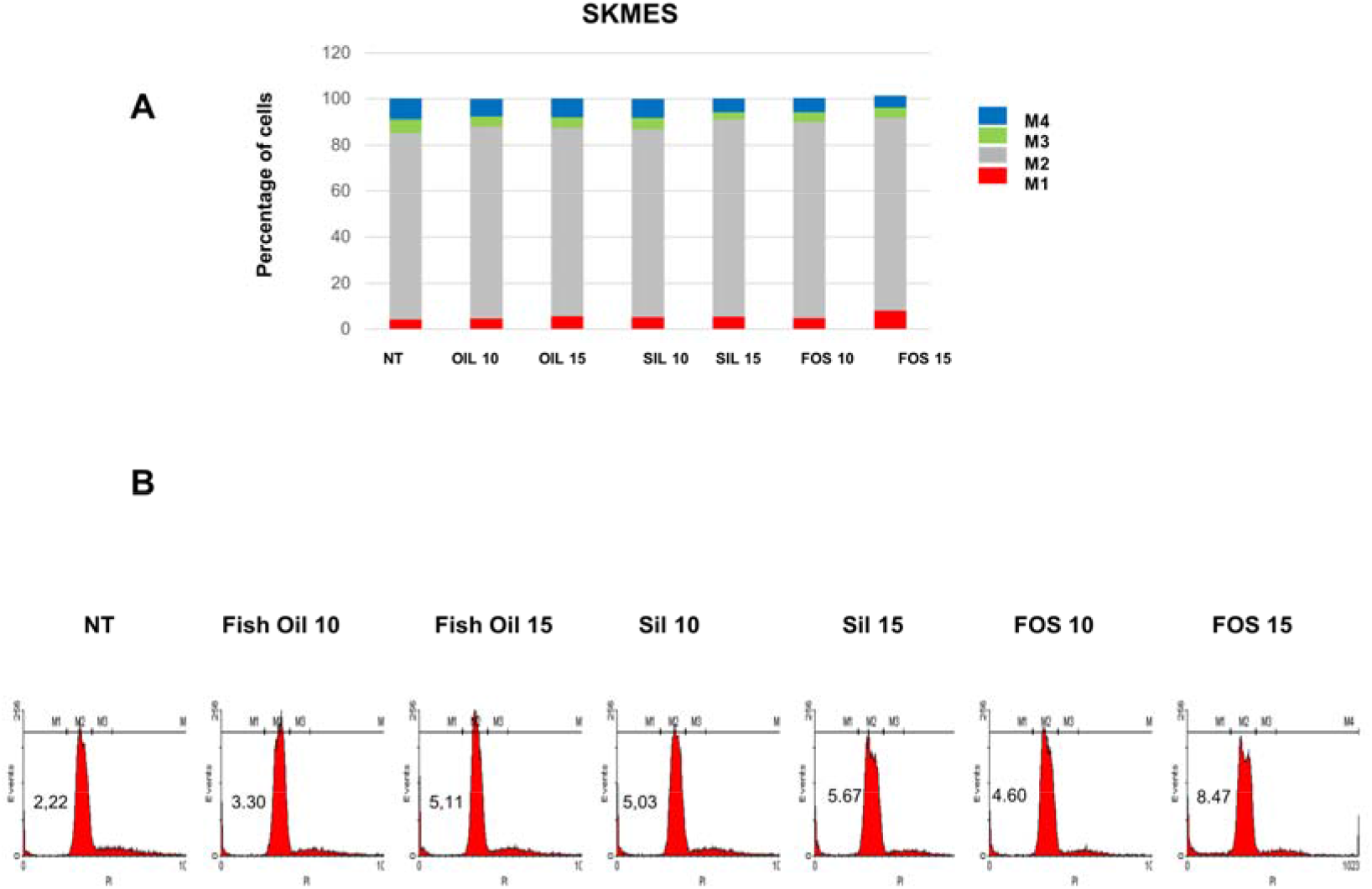
Effect of fish oil, silica and FOS on cell cycle events in SKMES. SKMES were cultured for 24 hours with fish oil (10 and 15 μg/ml), silica (10 and 15 μg/ml) and fish oil in silica (FOS) (10 and 15 μg/ml) and cell cycle was assessed by flow cytometry. **A.** Data expressed as percentage of cells (mean±SD). * p<0.05. **B.** Representative histograms are shown.

### 3.3 Effects of fish oil, silica and FOS on colony formation ability in NSCLC cell lines

In A549 (Figure 5A), in Colo699 (Figure 5B) and in SKMES (Figure 5C) only FOS at 10 μg/ml significantly reduced colony number when compared to untreated cells or to the other experimental conditions, including fish oil alone at the same concentration.

**Figure 5.**
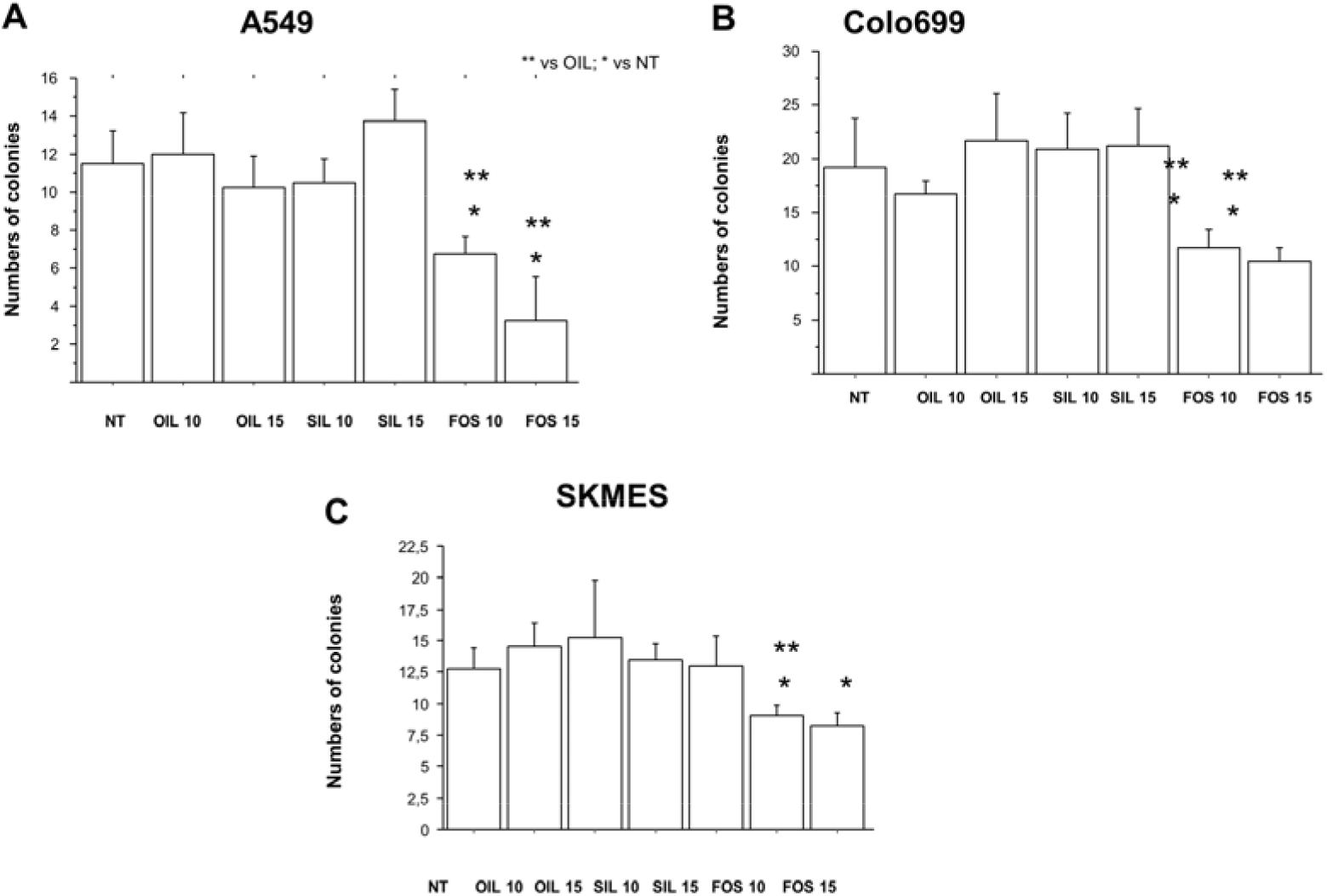
Effect of fish oil, silica and FOS on colony formation ability in NSCLC cell lines. NSCLC cell lines, A549 (**A**), Colo699 (**B**), and SKMES (**C**) were cultured for 24 hours with fish oil (10 and 15 μg/ml), silica (10 and 15 μg/ml) and fish oil in silica (FOS) (10 and 15 μg/ml)and colony forming ability was assessed by clonogenic assay. Data expressed as percentage of control (NT) (mean±SD). * p<0.05.

### 3.4 Effects of fish oil, silica and FOS on mitochondrial stress in NSCLC cell lines

Mitochondria play an essential role in the survival and maintenance of cancer cells and have been considered a target for anticancer agents.^22^ In A549, mitochondrial stress was increased upon exposure to fish oil, to silica and to FOS at both the tested concentrations. Both concentrations of FOS exerted a significant stronger activity on mitochondrial stress than fish oil or silica alone (Figures 6A-6B). In Colo699, mitochondrial stress was increased upon exposure to silica at 15 μg/ml and to FOS at both the tested concentrations. Both concentrations of FOS exerted a significant stronger activity on mitochondrial stress than fish oil or silica alone at both the tested concentrations (Figures 7A-7B).

**Figure 6.**
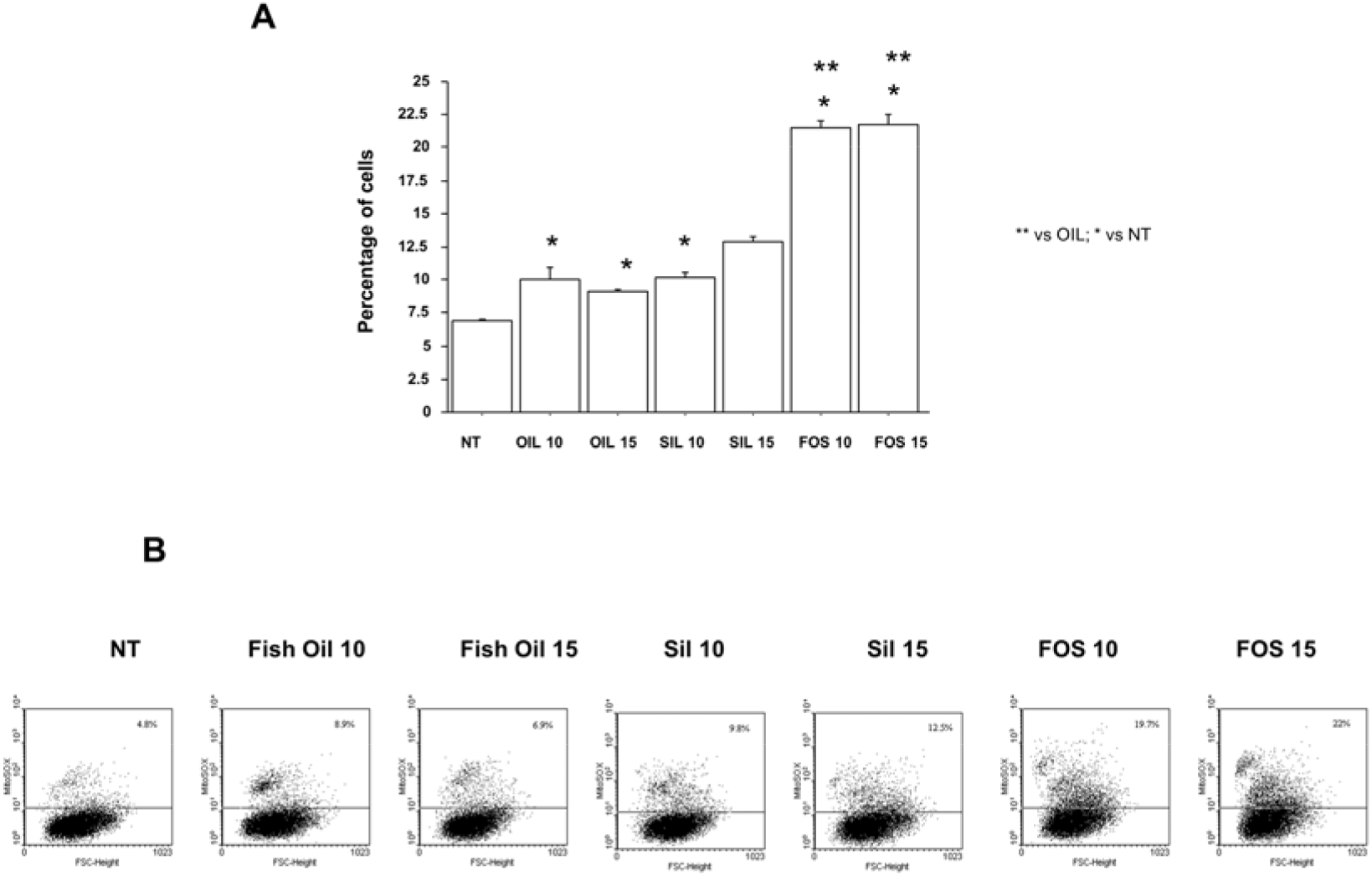
Effect of fish oil, silica and FOS on mitochondrial superoxide production in A549 cells. A549 cells were cultured for 24 hours with fish oil (10 and 15 μg/ml), silica (10 and 15 μg/ml) and fish oil in silica (FOS) (10 and 15 μg/ml) and mitochondrial superoxide production was assessed using MITOSOX by flow cytometry. Data expressed as percentage of positive cells (mean±SD). * p<0.05. **B.** Representative dot plots are shown.

**Figure 7.**
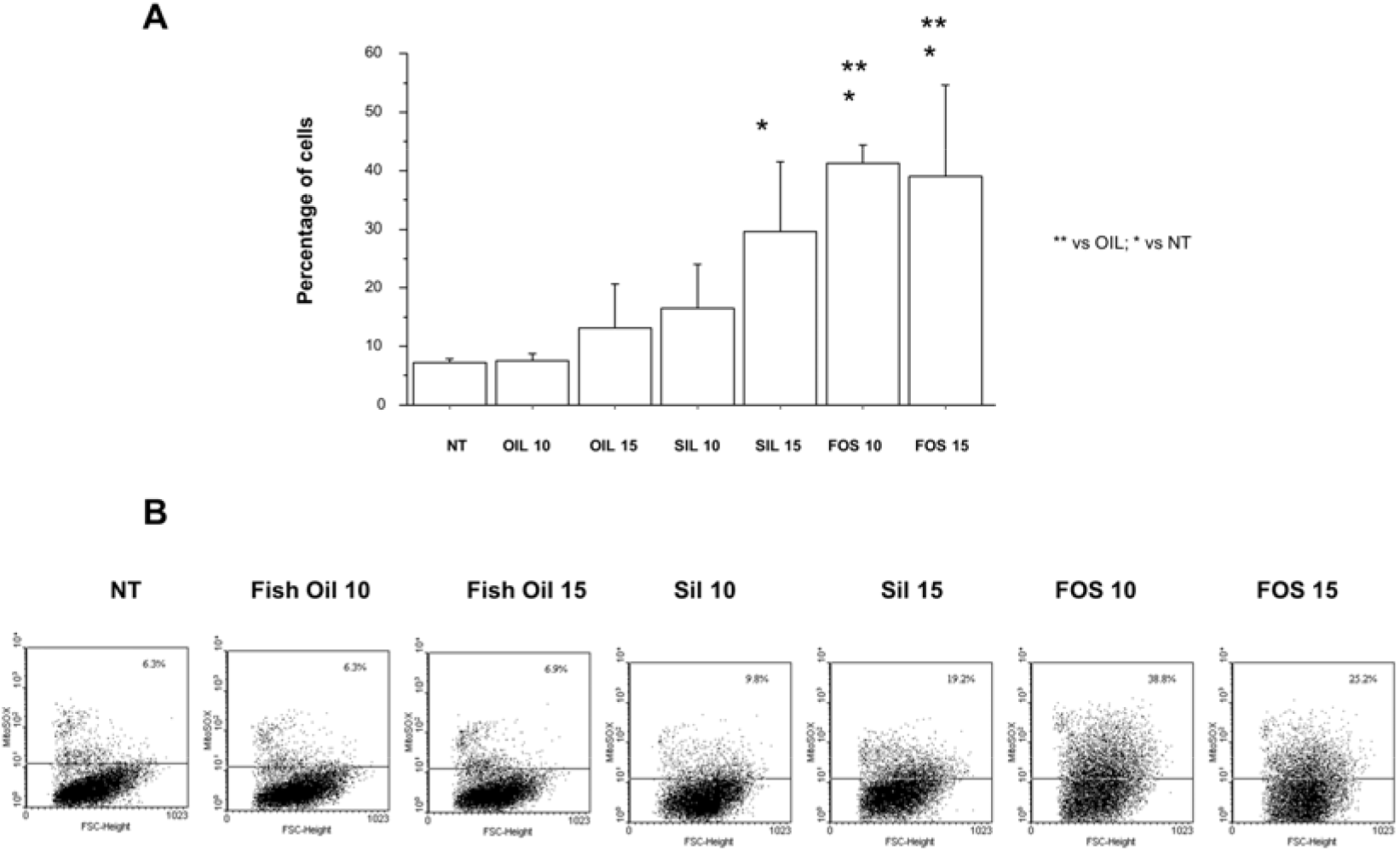
Effect of fish oil, silica and FOS on mitochondrial superoxide production in Colo699 cells. Colo699 cells were cultured for 24 hours with fish oil (10 and 15 μg/ml), silica (10 and 15 μg/ml) and fish oil in silica (FOS) (10 and 15 μg/ml) and mitochondrial superoxide production was assessed using MITOSOX by flow cytometry. Data expressed as percentage of positive cells (mean±SD). * p<0.05. **B.** Representative dot plots are shown.

In SKMES, mitochondrial stress was increased upon exposure to fish oil at 10 μg/ml and to silica and to FOS at both the tested concentrations. Both concentrations of FOS exerted a significant stronger activity on mitochondrial stress than fish oil alone at both the tested concentrations (Figures 8A-8B).

**Figure 8.**
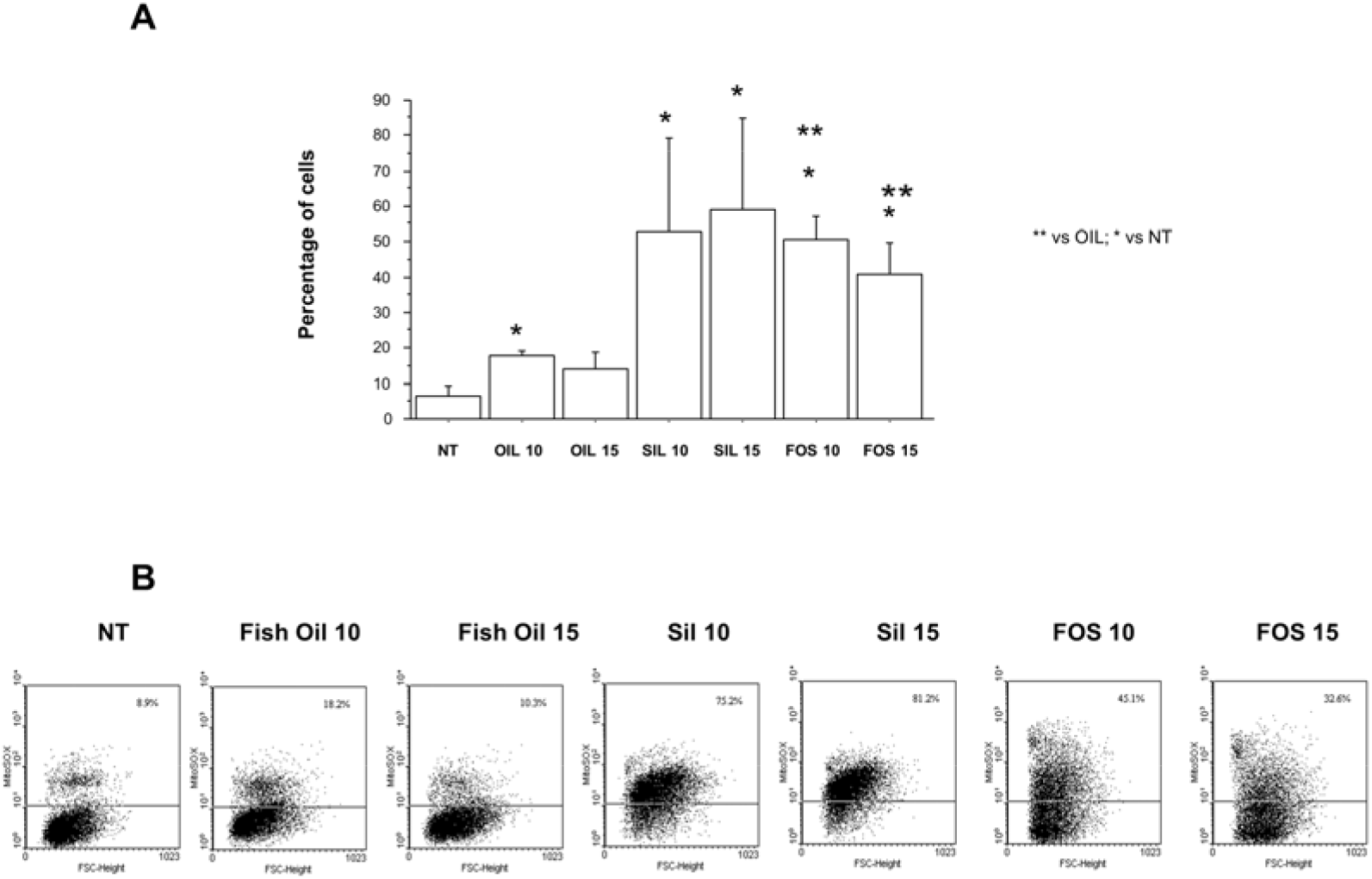
Effect of fish oil, silica and FOS on mitochondrial superoxide production in SKMES cells. SKMES cells were cultured for 24 hours with fish oil (10 and 15 μg/ml), silica (10 and 15 μg/ml) and fish oil in silica (FOS) (10 and 15 μg/ml) and mitochondrial superoxide production was assessed using MITOSOX by flow cytometry. Data expressed as percentage of positive cells (mean±SD). * p<0.05. **B.** Representative dot plots are shown.

### 3.5 Effects of fish oil, silica and FOS on ROS content in NSCLC cell lines

Upregulation of mitochondrial superoxide leads to increased intracellular ROS levels.^23^ In A549, it was observed a significant increase in ROS levels upon exposure to fish oil at both the tested concentrations, to silica 15 μg/ml and to both concentrations of FOS. Both concentrations of FOS exerted a significant stronger activity on ROS than fish oil alone at the same concentration (Figures 9A-9B).

**Figure 9.**
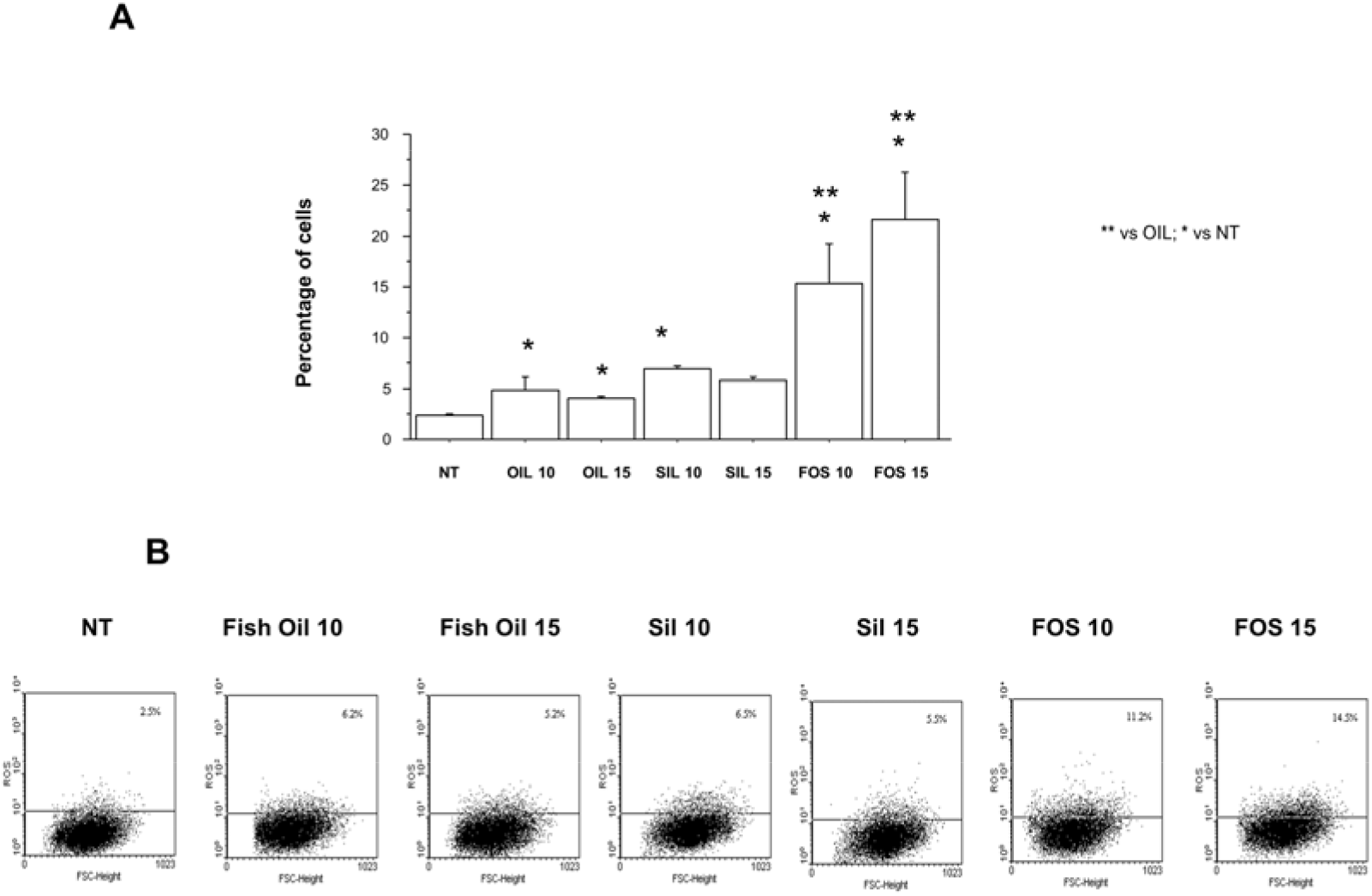
Effects of fish oil, silica and FOS on ROS production in A549 cells. A549 were cultured for 24 hours with fish oil (10 and 15 μg/ml), silica (10 and 15 μg/ml) and fish oil in silica (FOS) (10 and 15 μg/ml) and ROS production was assessed using DCFH-DA by flow cytometry. Data are expressed as percentage of positive cells (mean±SD). * p<0.05. **B.** Representative dot plots are shown.

In Colo699, it was observed a significant increase in ROS levels upon exposure to fish oil and to FOS at both the tested concentrations. FOS at 10 μg/ml exerted a significant stronger activity on ROS than fish oil alone at the same concentration (Figures 10A-10B). In SKMES, it was observed a significant increase in ROS levels upon exposure to FOS at 10 μg/ml and FOS at 10 μg/ml exerted a significant stronger activity on ROS than fish oil or silica alone at the same concentration (Figures 11A-11B).

**Figure 10.**
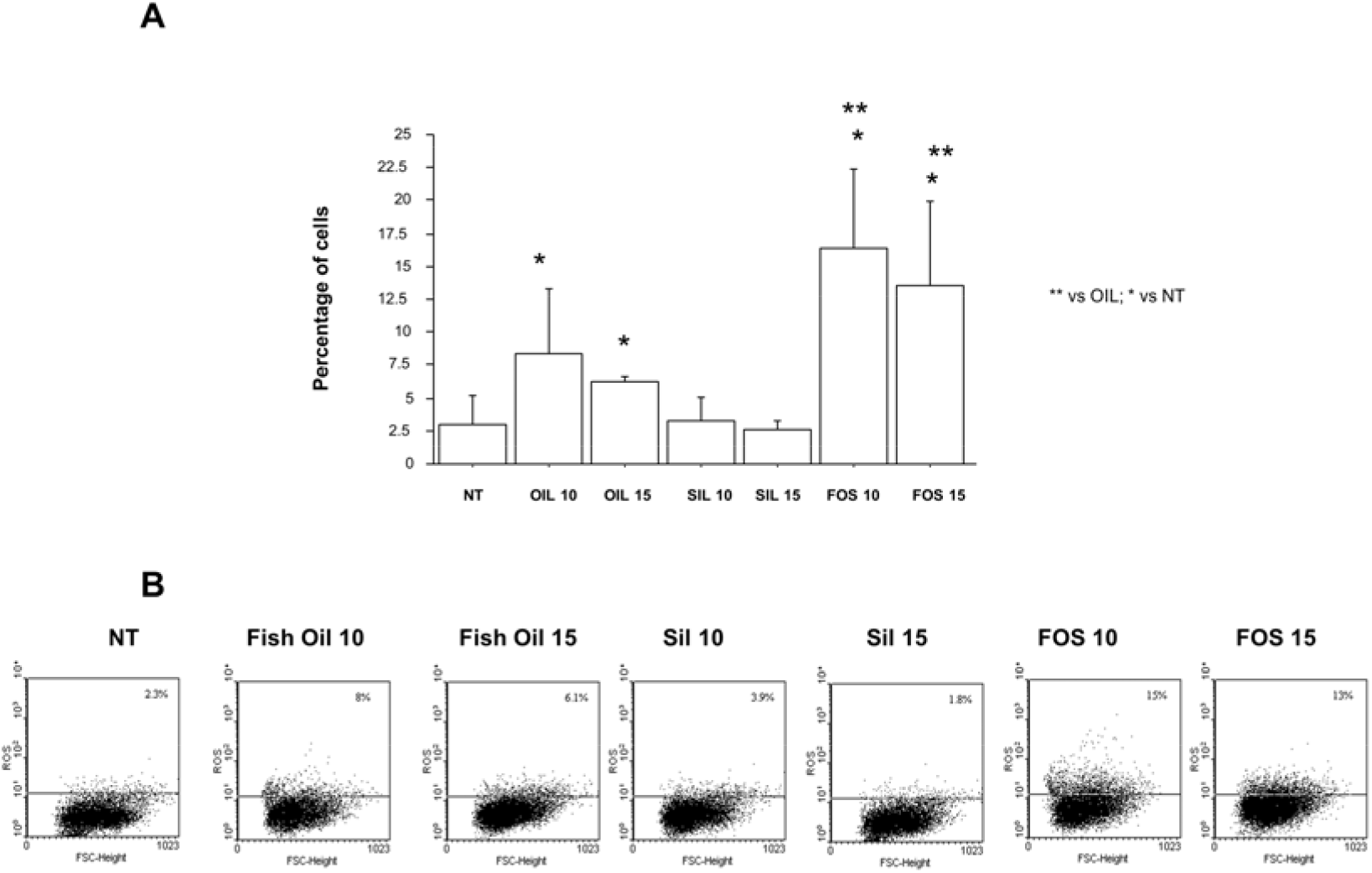
Effects of fish oil, silica and FOS on ROS production in Colo699 cells. Colo699 were cultured for 24 hours with fish oil (10 and 15 μg/ml), silica (10 and 15 μg/ml) and fish oil in silica (FOS) (10 and 15 μg/ml) and ROS production was assessed using DCFH-DA by flow cytometry. Data expressed as percentage of positive cells (mean±SD). * p<0.05. **B.** Representative dot plots are shown.

**Figure 11.**
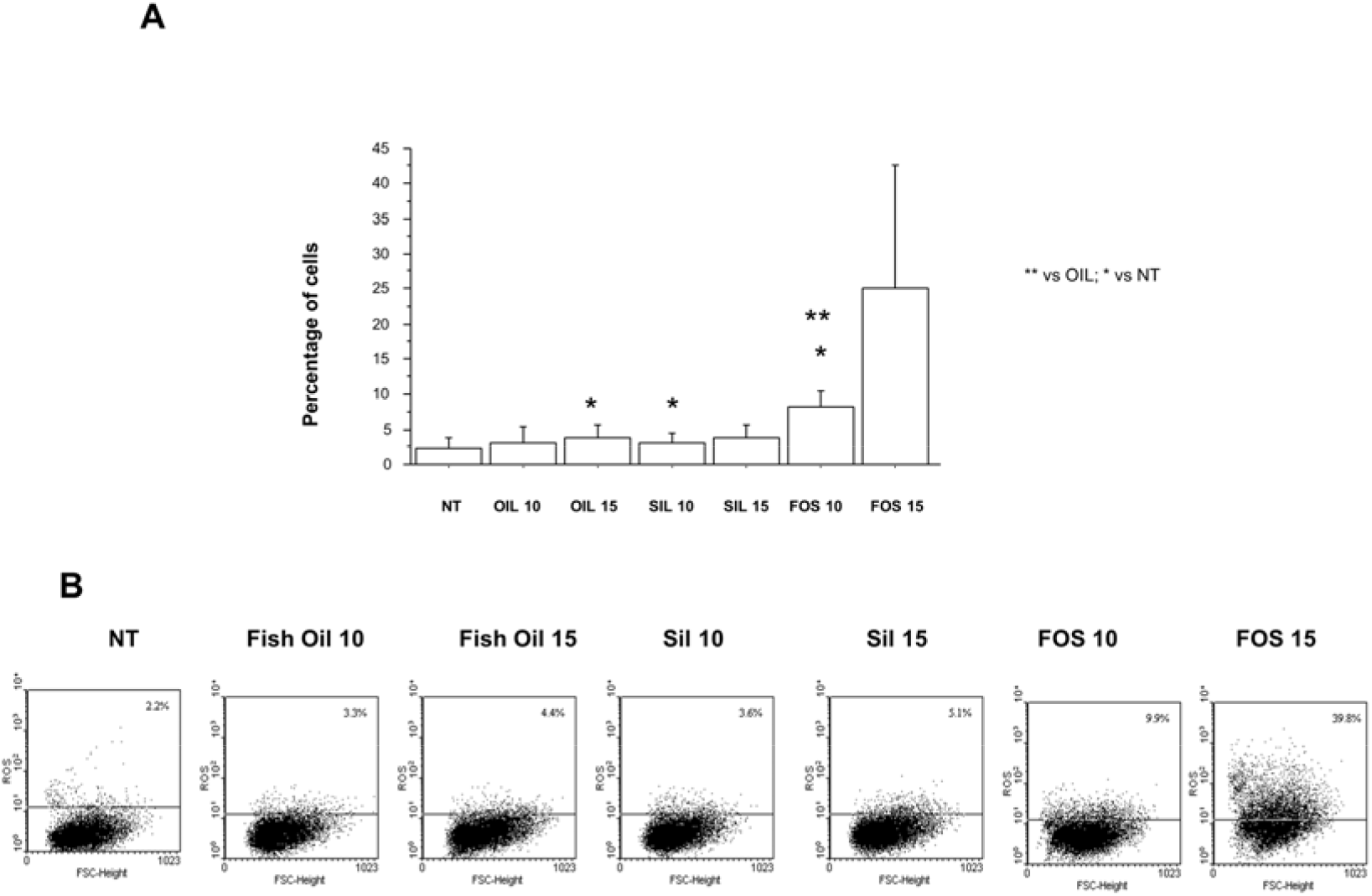
Effects of fish oil, silica and FOS on ROS production in SKMES cells. SKMES were cultured for 24 hours with fish oil (10 and 15 μg/ml), silica (10 and 15 μg/ml) and fish oil in silica (FOS) (10 and 15 μg/ml) and ROS production was assessed using DCFH-DA by flow cytometry. Data expressed as percentage of positive cells (mean±SD). * p<0.05. **B.** Representative dot plots are shown.

## 4. Discussion

The use of whole fish oil alone or encapsulated in silica particles in the present study represents an usual approach since the majority of the *in vitro* studies on the anti-cancer effects of omega-3 PUFAs assess the effects of the two major single EPA and DHA components in non esterified, acid form.^24^

The toxic effects of specific compounds on cancer cells can be evaluated by anlyzing early events looking at cell cycle alterations or late events assessing colony forming ability. Data reported herein demonstrate that FOS altered cell cycle phases in all the tested cell lines. At ultralow concentration of 10 μg/ml FOS was able to increase the number of apoptotic cells (identified as cell with reduced DNA content) in adenocarcinoma cell lines and to reduce colony forming ability in all the tested cancer cell lines. FOS was more effective than *AnchoisOil* fish oil alone at the same concentration.

Single components of fish oil, EPA and DHA, are able to induce toxic effects (cell apoptosis o reduced cell proliferation) on lung cancer cells, but at higher concentrations ranging from 50 to 75 μg/ml.^25^ These findings further support the effectiveness of this new silica-based formulation comprised of whole fish oil heterogenized in mesoporous silica. Hovewer, FOS at both tested concentrations does not increase cell apoptosis but reduces colony forming ability in squamous carcinoma cell line (SKMES). Taken togeheter, these data suggest that adenocarcinoma cells arising from distal airway epithelium are more sensitive in terms of early events (cell apoptosis) than cancer epithelial cells from the proximal airways. Future studies are needed to validate further higher concentrations of FOS for inducing early cell apoptosis also in lung squamous carcinoma.

With regard to late anti-cancer events, FOS reduced the colony forming ability in all the tested cancer cells. This could be related to the alteration on arachidonic acid (AA) metabolism. Indeed, the increased concentration of omega-3 PUFAs in cancer cells, associated with a statistically significant reduction in AA concentrations, suppresses AA-derived eicosanoid (PGs) biosynthesis thereby decreasing PGE_2_ concentration.^26^ As mentioned above, a further mechanism employed by omega-3 lipids to decrease PGE_2_ levels is linked to an inhibitory effect on COX-2, the cyclooxygenase enzyme responsible for the PGE_2_ synthesis.^27,28^ Prostaglandin E2 promotes tumor growth my multiple autocrine and paracrine mechanisms that increase cell proliferation and tissue invasion reducing cancer cell apoptosis and the efficiency of anti-cancer immune responses.^29^ In this context, it has been demonstrated that PGE contained in exudative pleural effusions from lung adenocarcinoma patients, is able to increase the colony forming ability of adenocarcinoma cell line (Colo699).^30^

Moreover, compounds capable of further increasing the oxidative stress within cancer cells have anticancer potential due to their ability to induce cell apoptosis, inhibiting cell proliferation and tissue invasiveness. Exerting a crucial role in ATP production and biosynthesis of macromolecules, mitochondria exert a relevant role in increasing intracellular ROS production. The mitochondrial metabolism is crucial for tumor proliferation, tumor survival, and metastasis.^22^ Compounds exerting antiproliferative and apoptotic activities in A549 human lung cancer alter mitochondrial function.^31^ Defects in mitochondrial ATP synthesis and mitochondrial membrane potential increase superoxide production thus leading to cell apoptosis. Multiple mechanisms may account for increased mitochondrial superoxide production. In this regard, inhibition of PIM kinases caused excessive mitochondrial fission and significant upregulation of mitochondrial superoxide.^32^

In addition, Bcl-2 levels are essential to mantain mitochondria integrity and preventing cell apoptosis. DHA downregulates Bcl-2 and upregulates Bax.^33^ When Bax transfers to the cell membrane, and binds to Bcl-2, it alters mitochodrial membrane potential and leads to increased superoxide production and to the release of cytochrome c, inducing cell apoptosis. Since FOS increase mitochondrial superoxide production and induce cell apoptosis, it is also conceivable that FOS could promote the interaction between Bax and Bcl-2. Moreover, fish oil can directly iincrease ROS levels. In this regard, it has been demonstrated that DHA increases ROS production by downregulating catalase.^32^

We ascribe the enhanced anticarcinogenic activity of *Omeg@Silica* in comparison to whole fish oil alone, to the hydrophilic nature of the embedding silica matrix and to large inner mesoporosity which allows to encapsulate within the hexagonal channels of periodic mesoporous silica an high load of bioactive *AnchoisOil*. Cancer cells are known to be acidic and are known to efficiently take up mesoporous silica nanoparticles functionalized with hydrophobic cancer drugs into the acidic organelles.^34^ Delivery of the omega-3 lipid molecules into the cancer cells leads to growth inhibition and cell impairment on all lung cancer‐cell lines examined. Yet, mesoporous SiO_2_ submicron particles are not toxic to human cells and end up preferentially localized in (acidic) lysosomes of cancer cells,^35^ which makes them ideally suited biocompatibile carriers for a host of hydrophobic anticancer drugs.

## 5. Conclusions

In this study we report the discovery of the high anticancer activity of a new material comprised of whole fish oil (*AnchoisOil*) extracted from anchovy fillet leftovers with biobased limonene sequestered within the inner mesoporosity of periodic mesoporous MCM-41 silica submicron particles. Tested *in vitro* on lung adenocarcinoma A549 and Colo699 cells, an aqueous formulation of these *Omeg@Silica* submicron particles in aqueous ethanol (FOS) is significantly more active than the fish oil alone in terms of effects on cell apoptosis, long term proliferation and mitochondrial superoxide ROS production. Less pronounced effects were observed when lung squamous cancer cells were treated with both fish oil and FOS.

Several aspects support further pre-clinical investigation of this new functional material in the treatment of lung cancer. First, the whole oil rich in omega-3 lipids, vitamin D_3_, zeaxanthin is directly obtained in natural form and at low cost from a freely available and abundant fishery by-product using a biobased and health-beneficial solvent (limonene) widely used in the food industry.^10^ No refinement of the resulting fish oil is needed to obtain the omega-3 lipids in diethylester form as it happens with most commercial omega-3 supplements.^14^

Second, the encapsulant material comprised of low polydisperse, submicron (269 nm) silica particles of large negative zeta potential (−32 mV) devoid of toxicity for healthy tissues^34,35^ is ideally suited to deliver the hydrophobic molecules comprising the *AnchoisOil*. These include EPA and DHA in natural (and highly bioavailable)^36^ form (DHA in position 2- and EPA in positions 1 and 3 of the triglyceride molecules) form as well as vitamin D_3_ and natural zeaxanthin. Both cholecalciferol^37^ and zeaxanthin^38^ are known to exert anticancer activity. This study, in conclusion, shows evidence that FOS exert anti-tumor effects on NSCLC affecting mitochondrial function and cell growth ability. Further studies to further characterize the molecular mechanisms originating the anti-cancer activity of FOS are needed to support the use of this new formulation in the therapy of lung cancer.

## Author Contributions

Caterina Di Sano and Claudia D’Anna conducted the largest number of experiments, participated in the interpretation of the data. contributed to write the manuscript and declare that they take the responsibility for the accuracy of the data analysis. Antonino Scurria and Claudia Lino extracted fish oil, synthesized mesoporous silica particles and incorporated fish oil in mesoporous silica affording the Omeg@Silica material. Mario Pagliaro revised the original manuscript and designed the Omeg@Silica structural investigation. Rosaria Ciriminna conceived the Omeg@Silica material and participated in the interpretation of the data. Elisabetta Pace designed the study, performed the statistical analysis of the data, contributed to the interpretation of the data, contributed to write the manuscript and declares that she has had access to and takes responsibility for the integrity of the data. All authors approved the final version of the manuscript.

## Data availability statement

Data used to support the findings of this study are included in the present article.

## Acknowledgments

We thank Agostino Recca Conserve Alimentari Srl (Sciacca, Italy) for kindly providing leftovers of anchovy fillets.

## Main abbreviations

NSCLC: Non-small cell lung cancer
SKMES: Squamous carcinoma cell line
PUFAs: Polyunsaturated fatty acids
DHA: Docosahexaenoic acid
EPA: Eicosapentaenoic acid
FOS: Fish oil encapsulated in mesoporous silica formulated in acqueous ethanol
DCFH-DA: Dichlorofluorescein diacetate
CTAB: Cetyltrimethylammonium bromide
AA: Arachidonic acid
ROS: Reactive oxygen species
PGE_2_: Prostaglandin E2
COX-2: Cyclooxygenase-2
NT: Non-treated

---

**Figure S1.**
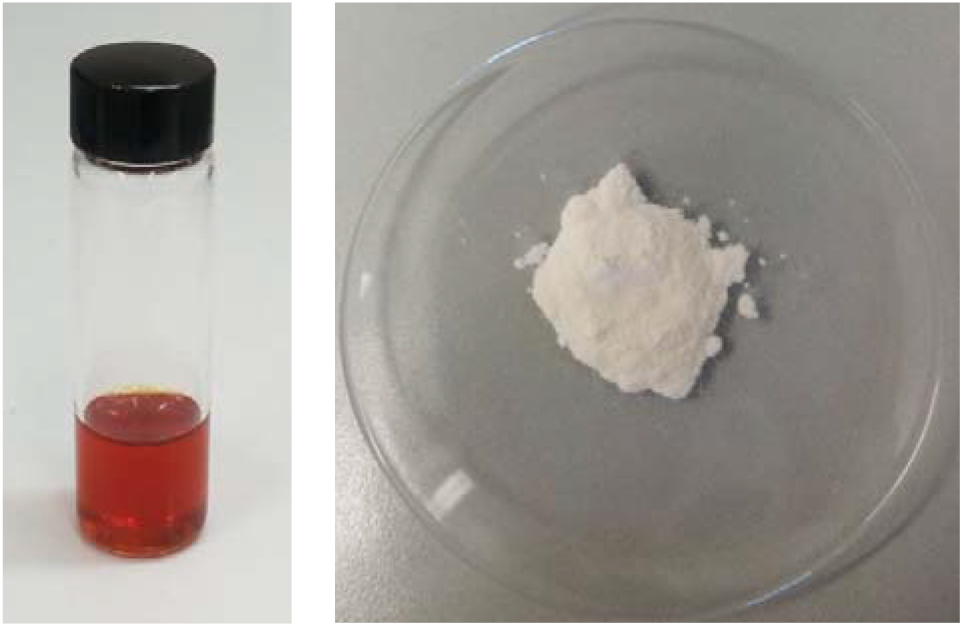
*AnchoisOil* extracted from anchovy filleting waste (*left*), *Omeg@Silica* comprised of MCM-41 silica loaded with 50% w/w fish oil (*right*).

